# Rice NIN-LIKE PROTEIN 4 is a master regulator of nitrogen use efficiency

**DOI:** 10.1101/2020.01.16.908558

**Authors:** Jie Wu, Zi-Sheng Zhang, Jing-Qiu Xia, Alamin Alfatih, Ying Song, Yi-Jie Huang, Guang-Yu Wan, Liang-Qi Sun, Hui Tang, Yang Liu, Shi-Mei Wang, Qi-Sheng Zhu, Peng Qin, Yu-Ping Wang, Shi-Gui Li, Chuan-Zao Mao, Gui-Quan Zhang, Chengcai Chu, Lin-Hui Yu, Cheng-Bin Xiang

## Abstract

Nitrogen (N) is one of the key essential macronutrients that affects rice growth and yield. Inorganic N fertilizers are excessively used to boost yield and generate serious collateral environmental pollution. Therefore, improving crop N use efficiency (NUE) is highly desirable and has been a major endeavor in crop improvement. However, only a few regulators have been identified that can be used to improve NUE in rice to date. Here we show that the NIN-like protein OsNLP4 significantly improves the rice NUE and yield. Field trials consistently showed that loss-of-*OsNLP4* dramatically reduced yield and NUE compared with wild type under different N regimes. In contrast, the *OsNLP4* overexpression lines remarkably increased yield by 30% and NUE by 47% under moderate N level compared with wild type. Transcriptomic analyses revealed that OsNLP4 orchestrates the expression of a majority of known N uptake, assimilation and signaling genes by directly binding to the nitrate-responsive *cis*-element in their promoters to regulate their expression. Moreover, overexpression of OsNLP4 can recover the phenotype of Arabidopsis *nlp7* mutant and enhance its biomass. Our results demonstrate that OsNLP4 is a master regulator of NUE in rice and sheds light on crop NUE improvement.

## Introduction

Nitrogen (N) is an essential macronutrient for the growth, development and production of plants (Crawford and Forde, 2002). Utilizable N resources in soil are very limited and therefore N fertilizer has been widely used to maintain high yield of crops in agriculture (Li et al., 2018; Lian et al., 2005). However, only a small fraction of the applied N is absorbed by plants, and a large portion is lost to the environment, causing serious environmental pollution (Garnett et al., 2009). N use efficiency (NUE) is defined as the total amount of yield in the form of grain or biomass achieved per unit of available N (Hawkesford, 2014). Improving NUE of crops is regarded as one of the most promising ways to solve the problem. However, the molecular mechanisms governing NUE is not well understood.

Plants have evolved effective mechanisms of N uptake and metabolism. Nitrate (NO_3_^-^) is the main N source absorbed by plants in well-aerated soil, which is mainly transported by nitrate transporter1 (NRT1), NRT2, chloride channel (CLC), and slow anion channel-associated homologues (SLAH) protein families (Krapp et al., 2014). After being uptaken into the cells, nitrate is reduced to ammonium (NH_4_^+^) by the action of nitrate and nitrite reductase (NR and NiR), then assimilated into amino acids by glutamine synthetase (GS) and glutamate synthase (GOGAT) (Hirel et al., 2011). In addition to its role as a major nutrient for plants, nitrate also acts as an important signaling molecule for numerous developmental processes (Ho and Tsay, 2010; Krouk et al., 2010a). Arabidopsis nitrate regulated1 (AtANR1) was the first identified nitrate regulator that functions in lateral root branching in response to nitrate (Zhang, 1998). AtNRT1.1 (CHL1), a dual-affinity transporter, acts as a NO_3_^-^ sensor that regulates the primary NO_3_^-^ response by changing the phosphorylation status of its threonine 101 residue (Park et al., 2009; Wang et al., 2009). The NO_3_^-^ inducible protein kinase calcineurin B-like-interacting protein kinases23 (AtCIPK23) and AtCIPK8 could up regulate or down regulate the expression of CHL1 respectively, under different nitrate concentrations (Hu et al., 2009; Park et al., 2009). Furthermore, nitrate regulatory gene2 (*AtNRG2*) was found to mediate nitrate signaling in *Arabidopsis* through modulating AtNRT1.1 expression and interaction with NIN-like protein7 (NLP7) (Xu et al., 2016). A few transcription factors (TFs) have been identified in the regulation of nitrate metabolism. For example, transcription factors lateral organ boundary domain 37/38/39 (AtLBD37/38/39) are negative regulators of NO_3_^-^ regulated genes (Rubin et al., 2009). Transcription factors squamosa promoter binding protein-like9 (AtSPL9), TGACG sequence-specific binding protein1 (AtTGA1), AtTGA4, auxin signaling F-box3 (AtAFB3), and NAC domain containing protein4 (AtNAC4) have also been identified as nitrate regulators involved in early nitrate response signaling (Alvarez et al., 2014; Krouk et al., 2010b; Vidal et al., 2014; Vidal et al., 2010).

Rice is one of the most important food crops in the world. N is a key factor affecting the growth and yield of rice. However, the molecular mechanisms underlying rice NUE is not well understood. The members of OsNRT1/2 family have received the most exploration and research to date. It has been reported that OsNRT1 encodes a constitutive component of the low-affinity nitrate uptake transporter (Lin et al., 2000). The variation in *OsNRT1*.*1B* largely explains nitrate-use divergence between *indica* and *japonica* and that *NRT1*.*1B*-*indica* can potentially improve the NUE of *japonica*. Moreover, NRT1.1B can interact with a phosphate signaling repressor SPX domain containing protein4 (SPX4) to coordinate utilization of N and phosphorus (Hu et al., 2019; Hu et al., 2015). Overexpression of *OsNRT1*.*1A* in rice significantly improves NUE and grain yield, and maturation time is also significantly shortened (Wang et al., 2018). Another nitrate transporter OsNPF6.1^HapB^ confers high NUE and increases yield under low nitrogen supply (Tang et al., 2019). Nitrate assimilation related2.1 (OsNAR2.1) regulates the balance of ammonium and nitrate by interacting with multiple NRT2 family members, including OsNRT2.1, OsNRT2.2 and OsNRT2.3a (Yan et al., 2011). In addition, other genes have been reported to affect N metabolism in plants, such as dense and erect panicle1 (DEP1), which interacts *in vivo* with both the Gα (RGA1) and Gβ (RGB1) subunits of the heterotrimeric G protein, and reduced RGA1 or enhanced RGB1 activity inhibits N responses (Huang et al., 2009; Sun et al., 2014). The balanced opposing activities and physical interactions of the rice growth-regulating factor 4 (OsGRF4) transcription factor and the growth inhibitor DELLA confer homeostatic co-regulation of growth and the metabolism of carbon (C) and N (Li et al., 2018). The indica *OsNR2* allele has superior grain yield and NUE than the japonica allele, in part via feed-forward interaction with *OsNRT1*.*1B* (Gao et al., 2019).

Of all the plants, only legume species have successfully evolved the capacity to deal with N limitation by developing a symbiotic interaction with rhizobium. The initiation of nodule development depends on a specific transcription factor, nodule inception protein (NIN) (Borisov et al., 2003; Schauser et al., 1999). Loss of NIN function hinders the infiltration of rhizobium and the nodule formation (Marsh et al., 2007). Interestingly, the NIN gene family is widely present in higher plants and algae, including species that do not fix gaseous N. *CreNIT2*, a *NIN-like protein* (*NLP*) gene in *Chlamydomonas*, acts as a central regulator required for nitrate signaling and assimilation. CreNIT2 binds the promoter of nitrate reductase gene in the presence of nitrate and to enhance its transcription (Camargo et al., 2007). Arabidopsis NLPs play a central role in N signaling (Konishi and Yanagisawa, 2013; Liu et al., 2017; Marchive et al., 2013). *AtNLP6* and *AtNLP7* function as the master regulators in the early nitrate response. Disruption of *AtNLP7* or suppression of *AtNLP6* results in a N-starved phenotype and impaired nitrate signaling (Castaings et al., 2009; Konishi and Yanagisawa, 2013). Constitutive expression of the maize *ZmNLP6* or *ZmNLP8* can complement the Arabidopsis *nlp7* mutant by restoring nitrate signaling and assimilation (Cao et al., 2017). AtNLP7 is regulated by nitrate via a nuclear retention mechanism, which is controlled by the phosphorylation of the conserved serine205 by Ca^2+^-sensor protein kinases10/30/32 (AtCPK10/30/32) in the N-terminal region of AtNLP7 (Liu et al., 2017). AtNLP8, which localizes in the nucleus constantly, can bind directly to the promoter of *CYP707A2* in a nitrate-dependent manner to reduce the ABA content in seeds and promote seed germination (Yan et al., 2016). Protein-protein interactions mediated by the PB1 domain were observed between a variety of combinations of different Arabidopsis NLPs (Konishi and Yanagisawa, 2019).

We previously reported that *AtNLP7* significantly improves plant growth by coordinately enhancing N and C assimilation (Yu et al., 2016). To continue the investigation on the roles of NLPs in NUE, we initiated research to analyze the function of rice NLPs in NUE. There are six NLP family members in rice (Chardin et al., 2014). However, their functions are still largely unknown. Here we report that OsNLP4, the closest homolog to Arabidopsis NLP7, plays a pivotal role in N uptake, assimilation and signaling. Genetic analyses and field trials showed that loss-of-*OsNLP4* reduced biomass, yield as well as NUE, while constitutive overexpression of *OsNLP4* significantly increased plant biomass, NUE and yield under different N conditions. OsNLP4 has a broad impact on the transcriptome responding to N, and can coordinate a majority of the genes related to N utilization and signaling. Our results indicate that OsNLP4 is a master regulator of NUE and a promising candidate for NUE improvement of crops.

## Results

### *OsNLP4* greatly influences vegetative growth under different N conditions

To investigate the function of *OsNLP4* in rice, we generated two knockout mutants (*japonica* variety Zhonghua11 background, ZH11) with CRISPR technology and transgenic lines overexpressing *OsNLP4* (OE) under the control of rice *ACTIN1* promoter, and obtained an additional T-DNA insertion mutant *nlp4-3* (*japonica* variety Dongjin background, DJ) (Fig. S1).

When grown in hydroponic culture with different concentrations of nitrate, the OE plants exhibited significant growth advantages, and the seedlings of the *nlp4* mutants exhibited obvious growth retardation compared with wild type (WT), especially under low nitrate conditions (0.02 mM, 0.2 mM) (Fig. 1A, Fig. S2). The biomass of WT, *nlp4*, and OE lines were significantly affected by OsNLP4 under different N concentrations. The knockout mutants showed the lowest biomass (fresh weight per plant), while the OE lines showed the highest (Fig. 1B). Both shoot and root biomass displayed similar differences to whole plant biomass among the genotypes (Fig. 1C and D). Taken together, these results demonstrate that OsNLP4 plays crucial roles in rice vegetative growth.

**Figure 1.**
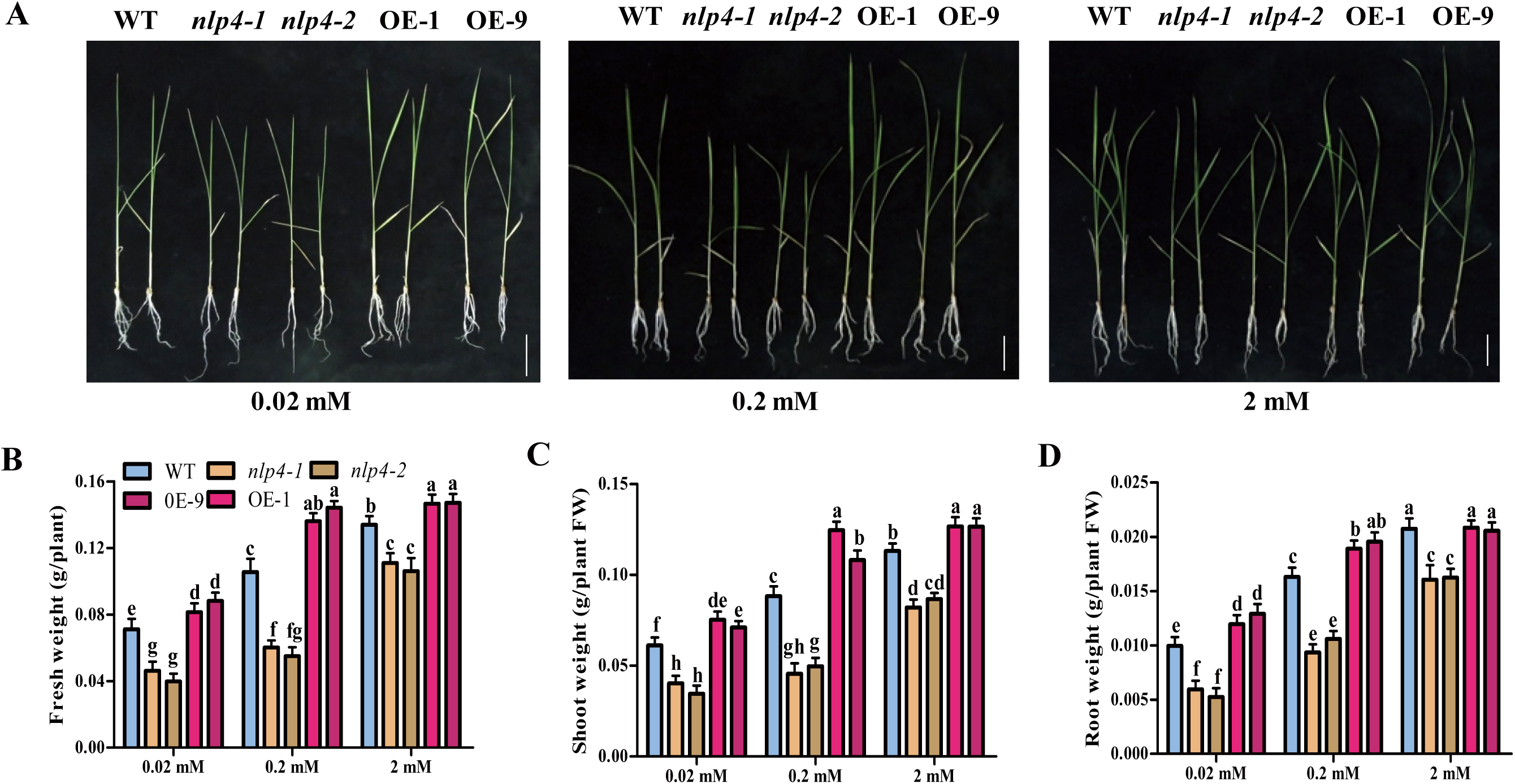
OsNLP4 influences plant growth under different N conditions. (A) Phenotypes of wild-type (WT), nlp4 mutants (nlp4-1/nlp4-2) and OsNLP4-OE lines (OE-1/OE-9) grown in hydroponic medium with different nitrate concentrations for 25 days. Bar = 4.0 cm. (B-D) Seedling biomass. Total fresh weight (B), shoot fresh weight (C), root fresh weight (D) of seedlings were measured. Values are the mean ± SD of four independent replications each containing 10 plants per genotype. P values are from the one-way ANOVA (The letters a, b and c indicate significant differences. P < 0.05). FW, fresh weight.

### OsNLP4 is a major determinant of rice NUE and crucial for yield

To investigate the role of OsNLP4 in rice NUE, we performed field trials for gain- and loss-of-*OsNLP4* genotypes in different years at two different locations in China: Chengdu and Lingshui. In Chengdu, the field trials were performed under low (LN), normal (NN) and high N (HN) conditions as described in Methods. A representative plant of each genotype grown under three N levels is shown in Fig. 2A. Grain yield per plant and per plot were tremendously affected by OsNLP4. The knockout mutants exhibited a significant decrease of yield per plant by an average of 29.8% under LN, 30.5% under NN, and 8.7% under HN compared with the wild type. In contrast, the OE lines showed a remarkable increase by an average of 38.0% under LN, 47.2% under NN, and 26.8% under HN (Fig. 2B). The same is true for actual yield per plot. The knockout mutants decreased yield per plot by an average of 29.5% under LN, 30.7% under NN, and 9.0% under HN, while the OE lines increased by an average of 37.4% under LN, 46.1% under NN, and 27.7% under HN compared with the wild type (Fig. 2C).

**Figure 2.**
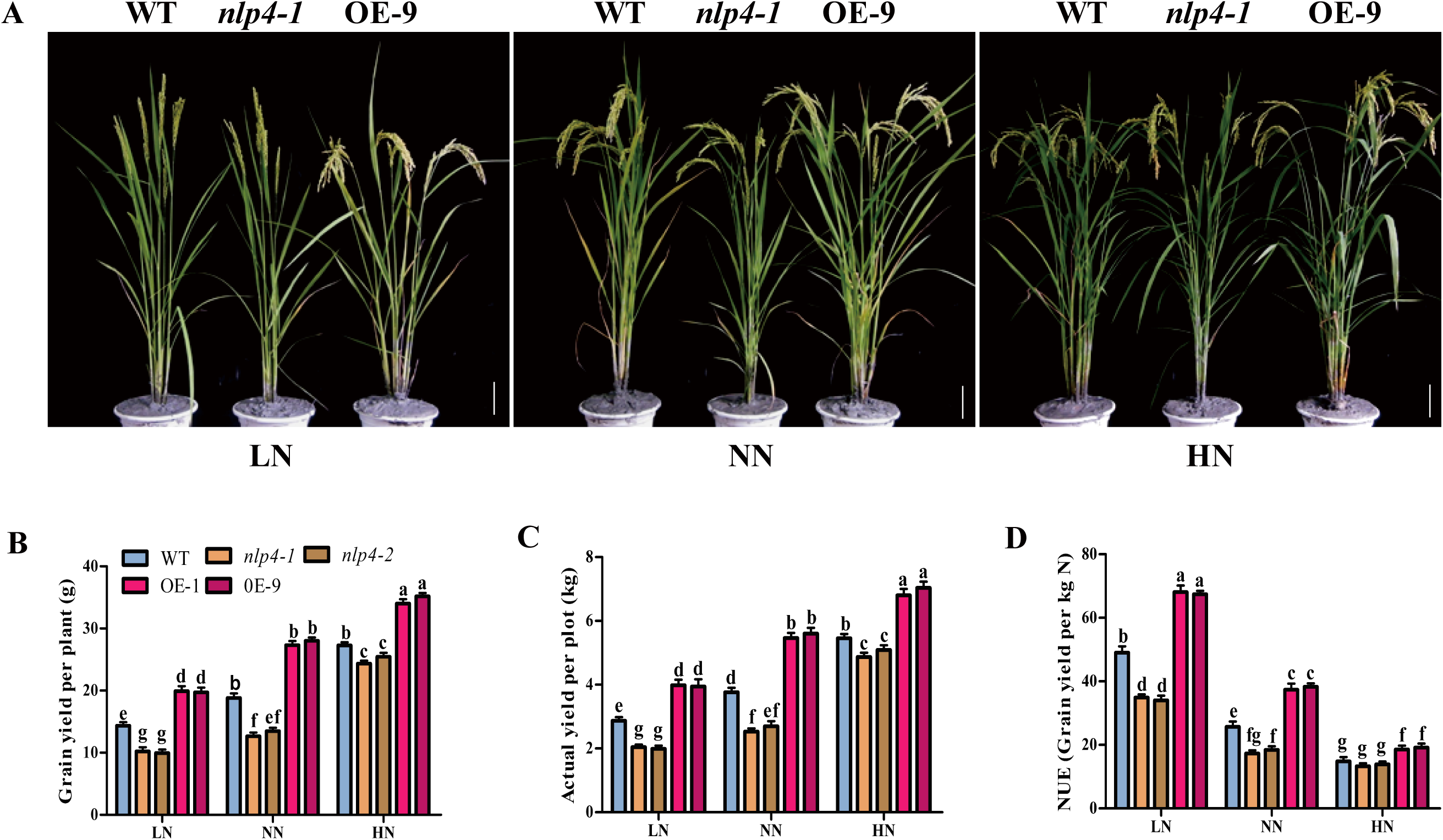
Field trials show OsNLP4 acting as a master regulator of NUE. (A) Growth phenotype of representative wild-type, nlp4 mutants (nlp4-1), and OsNLP4-OE plants (OE-9) at maturing stage in the field trial at Chengdu in 2019. LN, Low N conditions. NN, Normal N conditions. HN, High N conditions. Bar = 10 cm. (B-D) Grain yield per plant (B), actual yield per plot (C), NUE (D) of WT, nlp4 mutants (nlp4-1/nlp4-2) and OsNLP4-OE (OE-1/OE-9) plants under low N (LN), normal N (NN), high N (HN) conditions in the field trial at Chengdu (2019). Values are the means ± SD (four replications each containing 18 plants for grain yield per plant, 200 plants for actual yield per plot, NUE) (The letters a, b and c indicate significant differences. P < 0.05).

Based on the above yield and the amount of N fertilizer applied, NUE was calculated and shown in Fig. 2D. The OE lines remarkably enhanced NUE by an average of 37.3% under LN, 48.1% under NN, and 24.5% under HN compares with the wild type. In contrast, the knockout mutants significantly decreased NUE by an average of 28.9% under LN, 31.2% under NN, and 10.1% under HN.

In another field trial with normal N level in Lingshui, Hainan Province, the impacts of OsNLP4 on yield and NUE were reproduced (Fig. S3). The OE lines also significantly increased grain yield by 25.1% (grain yield per plant), 23.9% (actual yield per plot), and NUE by 24.3% (Fig. S3C). The T-DNA insertion mutant *osnlp4-3* showed dramatically decreased grain yield and NUE (Fig. S3D). Taken together, these results demonstrate that OsNLP4 is a major determinant of NUE and crucial for yield in rice.

### OsNLP4 is a master regulator coordinating NUE-related gene expression

To investigate the molecular mechanism by which OsNLP4 controls NUE and yield, we explore the genome-wide transcriptional landscape controlled by OsNLP4 in the response to N availability by profiling the transcripts of wild type (WT), knockout mutant *nlp4-1* (ko) and OE plants during a long-term hydroponic experiment with two N levels (LN with 0.02 mM nitrate and HN with 2 mM nitrate).

Transcriptomic differences between WT, ko and OE were determined by performing pairwise comparisons (WT vs. ko-LN, WT vs. ko-HN, WT vs. OE-LN and WT vs. OE-HN) of gene expression levels using the aligned reads so as to identify different expressed genes (DEGs) in each pair. Compared with the WT, DEGs with knockout or overexpression of *OsNLP4* were significantly different, especially between WT and ko under HN condition, indicating that OsNLP4 has a profound and broad influence on the transcriptome in response to nitrate (Fig. 3A and B, Fig. S4A).

**Figure 3.**
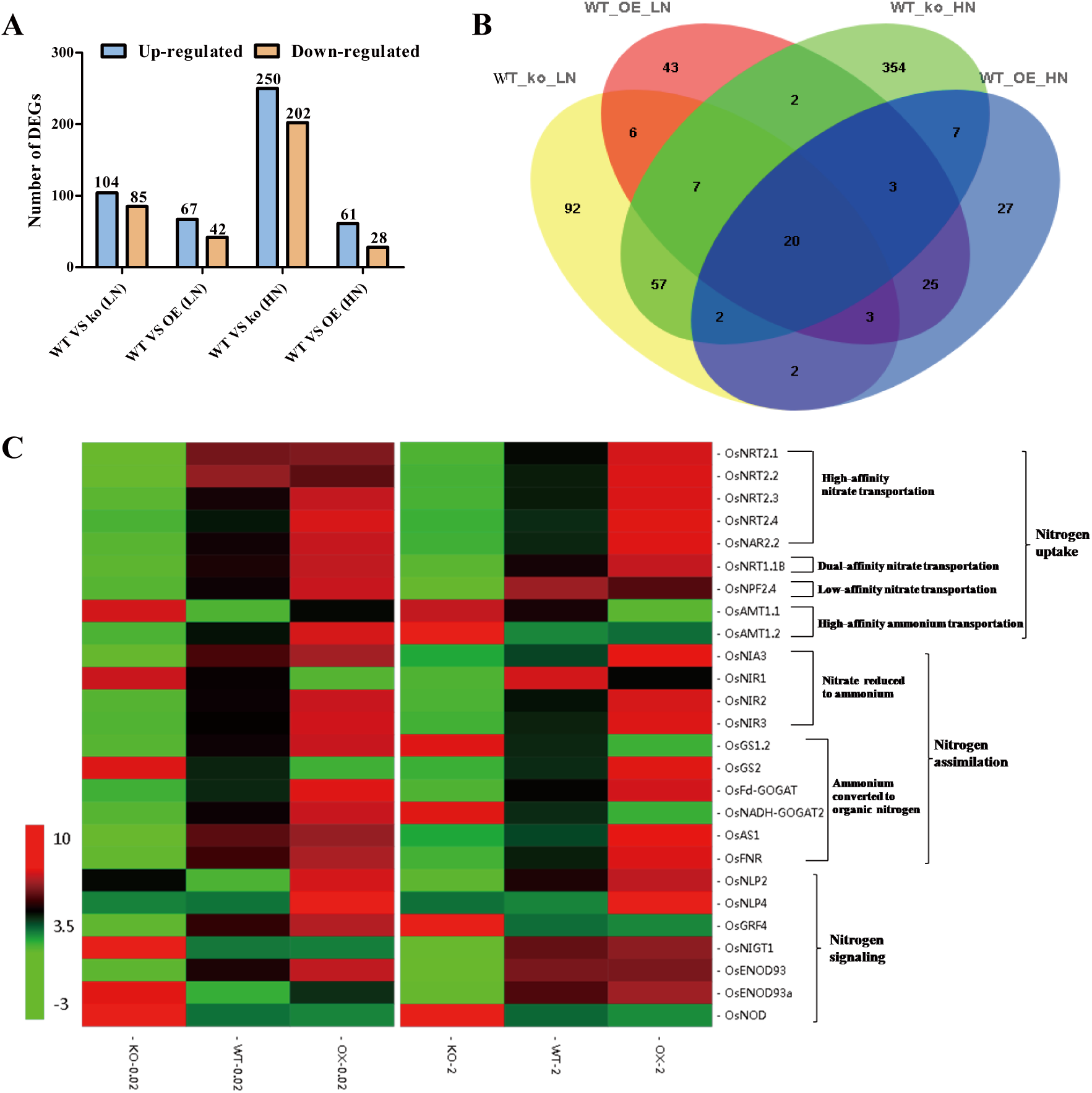
Expression profiling analysis of rice seedlings in different nitrate concentrations. (A) Number of up-regulated and down-regulated genes in WT vs. ko-LN, WT vs. ko-HN, WT vs. OE-LN and WT vs. OE-HN, as revealed by RNA-seq. (B) Venn diagrams analysis of the common and specific genes unique and shared between the different pair-wise comparisons. (C) Heatmap of the enriched NUE-related genes in DEGs in LN and HN conditions. Genes involved in nitro-gen uptake (including high-affinity nitrate transportation, dual-affinity nitrate transportation, low-affinity nitrate transportation and high-affinity ammonium transportation), assimilation (including nitrate assimilation and ammonium assimilation) and signal transduction were shown. The results were presented as log2 ratios. Only results with a P value < 0.05 and that have been confirmed in three independent experiments were included. A color code was used to visualize the data. LN (left figure): 0.02 mM KNO3; HN (right figure): 2 mM KNO3.

We further classified the DEGs based on Gene Ontology and KEGG pathway. Remarkably, genes involved in N uptake and metabolism were highly enriched in DEGs (Fig. S4B-C, Table S1), indicating that OsNLP4 may coordinately regulates the key genes in N utilization. Interestingly, as showed in Figure 3C, OsNLP4 appeared to have a preference to positively regulate the expression of high affinity nitrate/ammonium transporters, such as *OsNTR2*.*1, OsNTR2*.*2, OsNTR2*.*3, OsNTR2*.*4, OsNAR2*.*2, OsAMT1*.*2*, dual-affinity nitrate, such as *OsNTR1*.*1B*. Besides transporter genes, key genes involved in nitrate reduction, ammonium assimilation were dramatically down-regulated in *nlp4-1* mutant while significantly up-regulated in the *OsNLP4*-OE line. Moreover, several N signaling genes, such as *OsNLP2, OsGRF4, NITRATE-INDUCIBLE GARP-TYPE TRANSCRIPTIONAL REPRESSOR1* (*OsNIGT1*), *EARLY NODE 93* (*OsENOD93), OsENOD93a* were found differently expressed in *OsNLP4* mutant and overexpression plants. In addition, our data show that OsNLP4 also affected the expression of some genes related to C metabolism (Fig. S4B-C, Table S2).

The expression pattern of the genes involved in N response and metabolism was verified by RT-qPCR, which was largely in agreement with the RNA-seq data (Fig. S5). Taken together, these results suggest that OsNLP4 is a master regulator that orchestrates the N metabolism and signaling in rice.

### OsNLP4 directly modulates the expression of N metabolism genes by binding the NRE in their promoters

Consistent with the above results, the promoter regions of the key N metabolism genes harbor at least one NRE-Like *cis*-element for NLP binding (Table S3). To evaluate whether OsNLP4 directly binds to the NREs, we performed chromatin immunoprecipitation (ChIP) quantitative PCR assays and confirmed in vivo association of OsNLP4 with NRE-containing promoter fragments from the N assimilation genes, including *OsNRT2*.*1, OsNRT2*.*2, OsNRT2*.*3, OsNRT2*.*4, OsNRT1*.*1B, OsNIA1, OsNIA3, OsAMT1*.*1* and *OsGRF4* (Fig. 4A-I). This result was further verified with Y1H assay (Fig. 4J). These data indicate that the OsNLP4 may directly regulate the transcription of N absorption and assimilation genes.

**Figure 4.**
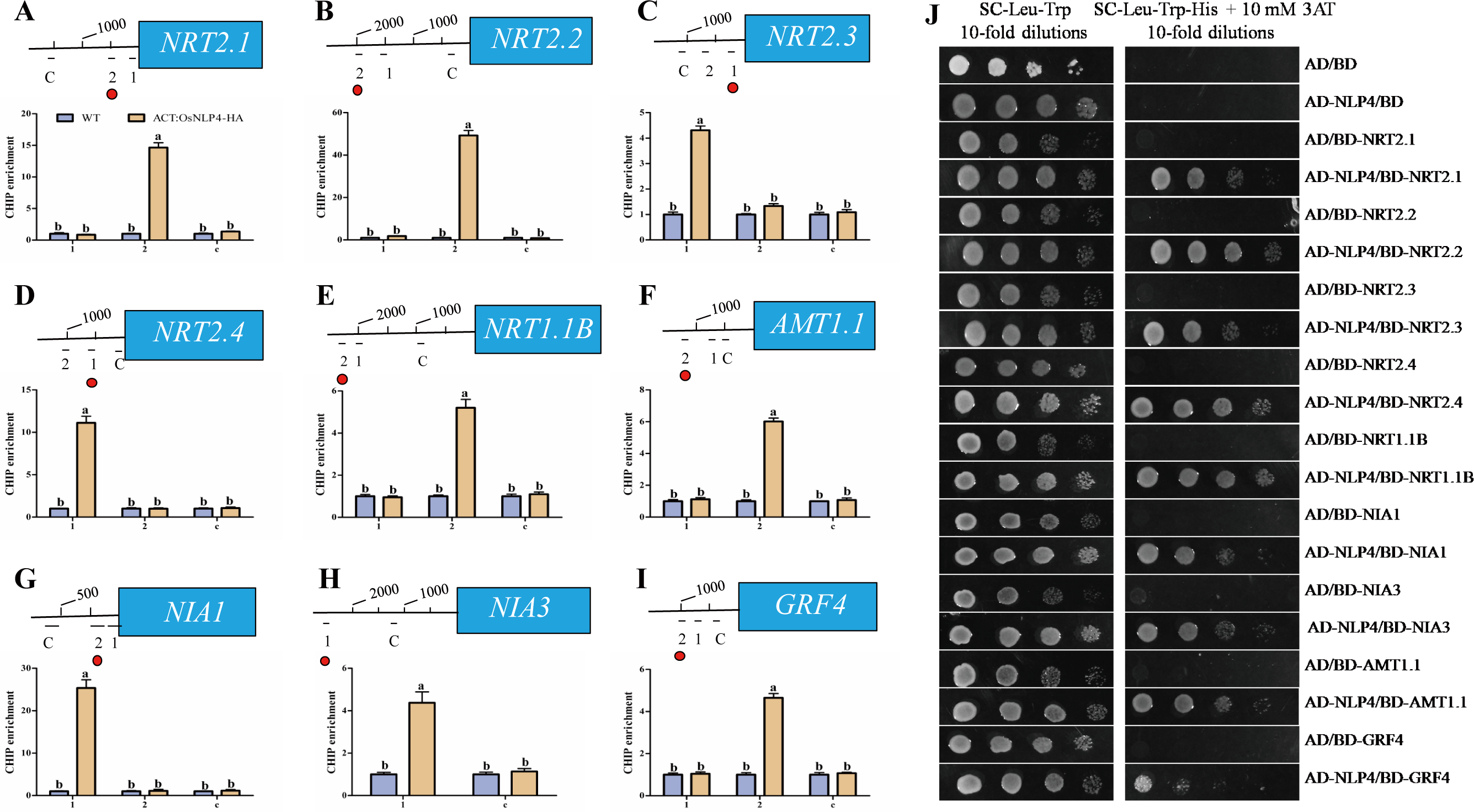
OsNLP4 binds to the promoters of many N-related genes directly. (A-I) OsNLP4 mediated ChIP–PCR enrichment (relative to input) of NRE-containing promoter fragments (marked with a dot) from (A) OsNRT2.1, (B) OsNRT2.2, (C) OsNRT2.3, (D) OsNRT2.4, (E) OsNRT1.1B, (F) OsNIA1, (G) OsNIA3, (H) OsAMT1.1 and (I) OsGRF4. Data are mean ± SD (n = 3). The letters a, b and c indicate significant differences (P < 0.05). (J) Y1H assay. Empty pAD and pBD were used as negative control. Serial yeast dilutions (1:1, 1:10, 1:100 and 1:1,000) were grown on different SD medium for 5 days.

### OsNLP4 directly modulates N uptake and assimilation

To confirm the above results of transcriptomic comparison, we investigated the role of OsNLP4 in regulating N absorption and assimilation. With a chlorate-sensitivity assay, we found that the *nlp4* mutants exhibited a much lower chlorate sensitivity while the OsNLP4-overexpressing lines showed significantly higher chlorate sensitivity than WT (Fig. 5A and B), demonstrating that OsNLP4 positively modulates nitrate uptake. We further demonstrated this by directly measuring ^15^N-nitrate uptake. The ^15^N accumulation in OE plants was significantly higher than that in WT while much lower in the knockout mutants (Fig. 5C). Moreover, total N and nitrate content were increased in OE lines but reduced in knockout mutants compared with those of the wild type (Fig. 5D and E).

**Figure 5.**
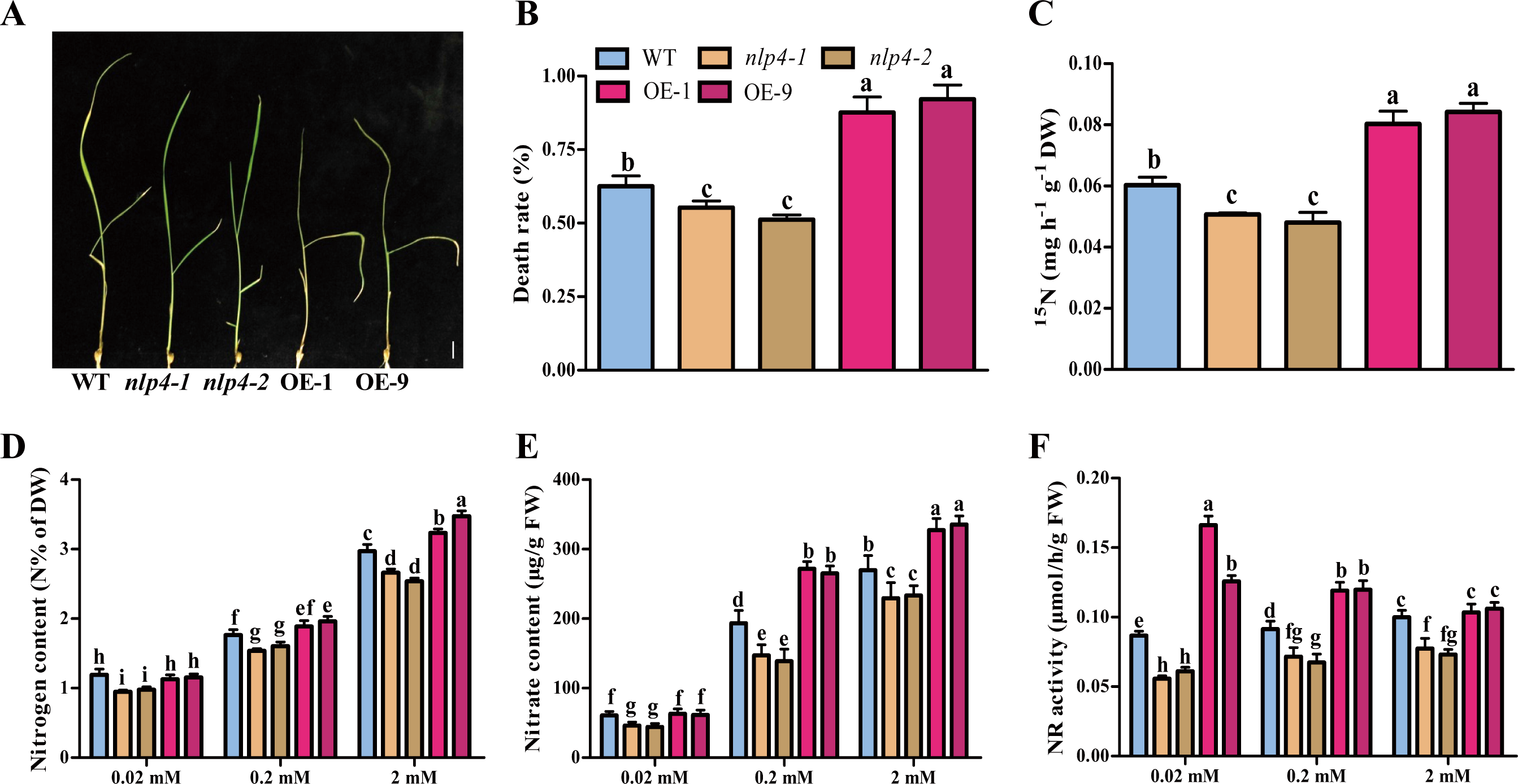
OsNLP4 regulates N uptake and assimilation. (**A**) Chlorate-sensitivity assay in wild-type, *nlp4* mutants and *OsNLP4*-OE plants treated with 2 mM chlorate. Bar = 1.0 cm. (**B**) Chlorate sensitivity was calculated by the mortality rate of plants poisoned by chlorate. Values are the mean ± SD of three replications each containing 50 plants per genotype (The letters a, b and c indicate significant differences. P < 0.05). (**C**) ^15^N accumulation assays in wild-type, *nlp4* mutants and *OsNLP4*-OE plants labeled with ^15^N-nitrate. Values are the mean ± SD of three replications each containing 15 plants per genotype (The letters a, b and c indicate significant differences. P < 0.05). (**D-F**) Contents of total N (D), and nitrate (E), enzyme activities of NR (F) in the plants grown under different nitrate conditions. 14-day-old seedlings grown in hydroponic medium with different concentrations of nitrate were used for metabolite analyses and enzymatic assays as described in Material and Methods. Values are the mean ± SD of four replications each containing 10 plants per genotype (The letters a, b and c indicate significant differences. P < 0.05). DW, dry weight. FW, fresh weight.

To confirm the role of OsNLP4 in regulating N assimilation, we investigated the N assimilation marker nitrate reductase and found that NR activity increased markedly in OE plants in contrast to the dramatic decrease in the *nlp4* mutants (Fig. 5F), indicating a higher nitrate assimilation efficiency in the *OsNLP4*-overexpressing plants, especially under low N conditions.

The above results were reproduced in an independent experiment with *nlp4-3* mutant (Fig. S6). Taken together, these data demonstrate that OsNLP4 regulates N uptake and assimilation.

### *OsNLP4* expression is responsive to N availability

To have a comprehensive understanding of *OsNLP4* expression in response to N availability, we made a detailed time-course analysis of *OsNLP4* expression. As shown in Figure 6A, when seedlings grown in normal N conditions (NN) were shifted to N starvation conditions (0N), *OsNLP4* transcript level was rapidly induced and plateaued with 5-fold increase after 2-hour N starvation, which was maintained as long as N starvation was imposed. When N was resupplied in the form of KNO_3_, *OsNLP4* transcript level was rapidly declined to its pre-induction level within 2 hours. However, when resupplied with NH_4_Cl, *OsNLP4* transcript level also decreased rapidly but maintained at a higher level compared with KNO_3_ treatment (Fig. 6A), while the KCl control treatment did not alter its transcript level. Moreover, *OsNLP4 promoter*::*GUS* transgenic plants exhibited similar response with strong glucuronidase (GUS) induction by N starvation and reversion by resupplied N (Fig. 6B).

**Figure 6.**
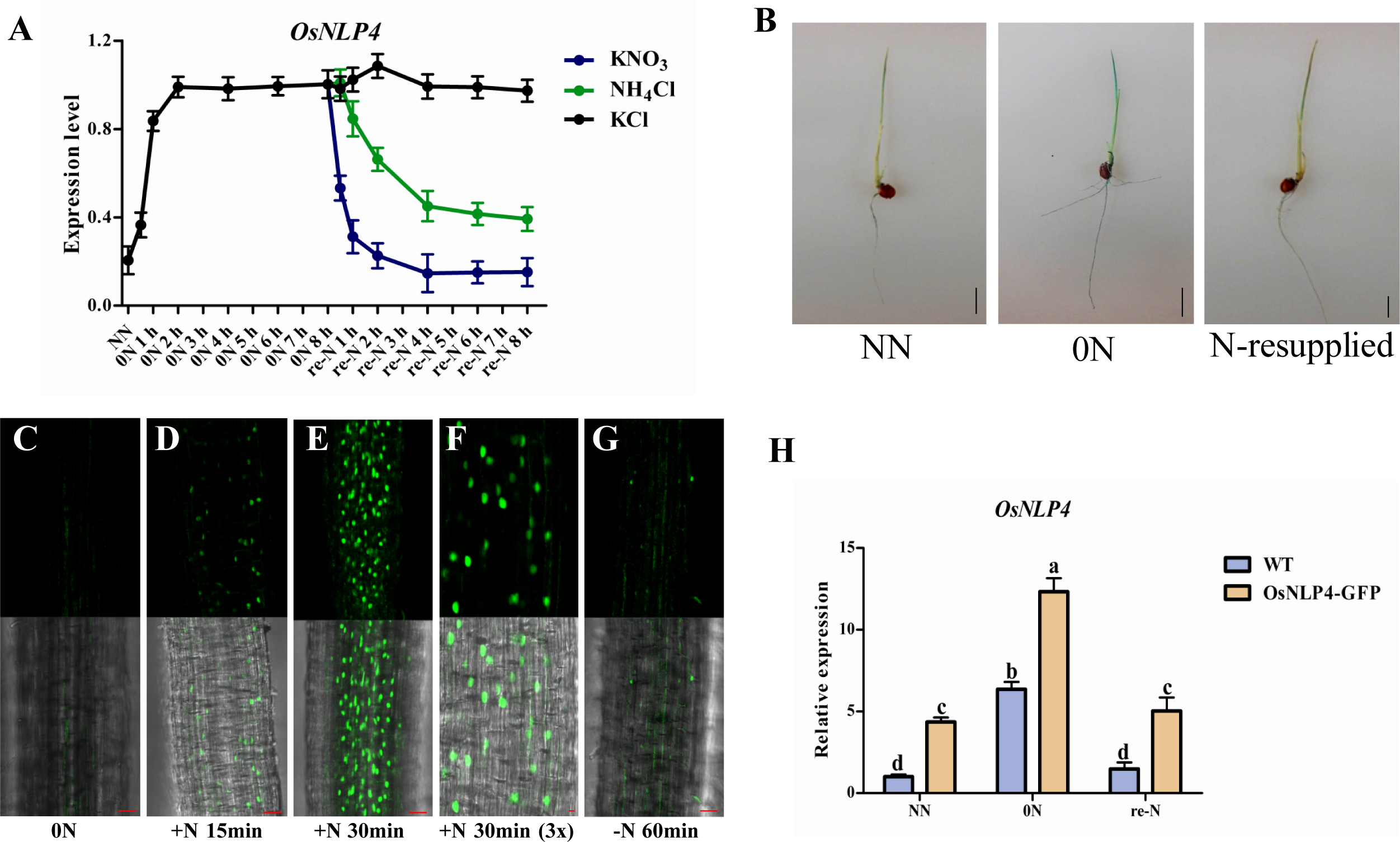
OsNLP4 expression is responsive to nitrogen availability. (A) Time-course analyses of OsNLP4 expression level in response to N availability. The time-course experiment with different nitrogen treatments (5 mM KCl, NH4Cl, and KNO3). RT-qPCR data are mean ± SD (n = 3). (B) GUS staining of 7-day-old OsNLP7promoter::GUS transgenic plants were cultured in different N conditions. NN: 2 mM KNO3; 0N: nitrogen free; N-resupply: nitrogen free for 6 days then with 2 mM KNO3 for 24h. Bar = 1.0 cm. (C-G) Subcellular localization of OsNLP4. (C) Seedlings grown on nitrogen-free medium. (D, E) N-starved seedlings were transferred to a nitrate medium (50 mM) and observed after 15 (D) and 30 min (E). (F) Partial enlargement of figure D in 30 times. (G) N-starved seedlings were transferred to a nitrate medium for 60 min, then to N-free medium again and observed after 60 min. Bar =50 μm. (H) Transcript levels of OsNLP4 in WT and OsNLP4 overexpression rice (ACTIN1::OsNLP4-GFP) under different N conditions. NN: 2 mM KNO3; 0N: nitrogen free; re-N: nitrogen free for 9 days then with 2 mM KNO3 for 24h. Values are the mean ± SD (n = 3). The letters a, b and c indicate significant differences (P < 0.05).

GUS activity was detected throughout the whole life of the reporter plant (Fig. S7A-K) with strong expression in coleoptile of germinating seeds, leaves and roots (Fig. S7A-D). Further observation showed that GUS activity was mainly detected in the epidermis and vascular tissues of leaves and roots (Fig. S7C-D and G), and also in stem, node, spikelet and anther (Fig. S7H-K). However, *OsNLP4* was not expressed in the root hairs (Fig. S7F). Consistent with the GUS staining result, qRT-PCR analyses of *OsNLP4* transcript level revealed a similar pattern with high expression in leaf and root (Fig. S7L).

To investigate the subcellular localization of OsNLP4 protein, we generated transgenic rice lines expressing the fusion of full length CDS of *OsNLP4* with GFP driven by the rice *ACTIN1* promoter. We checked the subcellular localization of OsNLP4 protein under N starvation and N re-addition conditions. OsNLP4 mainly localized in cytosol under N starvation (Fig. 6C). However, when nitrate was resupplied for 15 minutes, GFP signals were detected in the nucleus (Fig. 6D), by 30 minutes GFP signals were dramatically increased in the nucleus (Fig. 6E and F). This nuclear accumulation was rapidly reversed when N was withdrawn for 60 minutes (Fig. 6G). The dramatic change of GFP signal intensity in response to N availability strongly suggests a tight regulation of *OsNLP4* mRNA translation by N because the transcript level of *OsNLP4* was high in the wild type and *OsNLP4-GFP* overexpression plants under N starvation condition, and decreased once nitrate was resupplied (Fig. 6H).

### OsNLP4 enhances N assimilation and growth of the transgenic Arabidopsis

In order to investigate whether OsNLP4 can similarly modulate N assimilation in different plant species, we generated Os*NLP4*-overexpressing transgenic Arabidopsis in *nlp7-1* background. We found overexpression of Os*NLP4* not only fully rescued the N-deficient phenotypes of *nlp7-1*, but also significantly increased plant biomass under different nitrate conditions (Fig. S8A, C). Moreover, the overexpression plants had stronger roots (Fig. S8B, D) and grew better with larger rosette leaves and higher biomass when grown in soil (Fig. S8E and F). In addition, the expression of the genes involved in N assimilation and signaling was significantly up-regulated in overexpression lines compared with that of *nlp7-1* and WT (Fig. S9). These results suggest that *OsNLP4* is functionally conserved, and thus has potential applications to improve N use and plant growth in both monocot and eudicot crops.

## Discussion

Plant growth and yield are usually limited by soil N availability in most agricultural cropping systems. N fertilizer overuse leads to low NUE and serious environmental problems (Vitousek et al., 2009). Improving plant NUE is an important, cost-effective approach to boost crop yields in intensive agricultural systems worldwide. Many genes, most of which are related to N uptake and primary assimilation, have been applied to increase N assimilation in various plants during the past decades. However, only very few of these studies have showed phenotypic effect on NUE or other growth parameters on the plant by individually ectopic expression of these genes, and some even showed negative pleiotropic effects, implying the notion of single-point rate-limiting regulation being oversimplified for improving NUE (Andrews et al., 2004; McAllister et al., 2012; Xu et al., 2012). Application of transcription factors might be a more efficient strategy for improving NUE, because they have the capacity to coordinately regulate the expression of a set of genes so as to improve plant growth and NUE by enhancing multiple steps cooperatively instead of single step in a metabolic pathway. In the present study, we show that the transcription factor OsNLP4 acts as a master regulator to simultaneously coordinate many processes in N utilization and signaling pathway to significantly improve rice yield and NUE under different N conditions.

Likely benefiting from the simultaneously coordinating the expression of N-metabolism genes (Fig. 3, Fig. S4 and 5), *OsNLP4*-overexpressing rice significantly increased nitrate uptake and N assimilation (Fig. 5 and Fig. S6), thus enhanced plant growth (Fig.1 and Fig. S2), grain yield and NUE (Fig. 2 and Fig. S3) compared with wild type under all N conditions. On the contrary, plant growth, grain yield and NUE of the *osnlp4* mutants are significantly impaired compared with wild type.

NUE increased by 37.3%-48.1% in *OsNLP4*-overexpressing lines, while decreased by 28.9%-31.2% in *osnlp4* mutants compared with wild type under LN and NN, respectively. However, under HN, there is no difference in NUE between wild type and the mutants, while OsNLP4-overexpressing lines shows a small but statistically significant increase compared with wild type (Fig. 2). It is noteworthy that the field trial showed that grain yield per plot of the OsNLP4-overexpressing lines under LN and NN almost the same as that of WT under NN and HN, respectively (Fig. 2A), which indicates, with the help of OsNLP4, farmers can harvest similar grain yield as wild type with a 50% reduction of N fertilizer usage, thereby reducing the production cost and environmental pollution.

As the closest homolog of *AtNLP7* in rice (Chardin et al., 2014), *OsNLP4* can fully recover the N-starvation phenotype of the Arabidopsis *nlp7-1* mutant (Fig. S8). OsNLP4 also orchestrates the N utilization and signaling, thus improve the NUE under both N-rich and deficiency conditions, as revealed by our transcriptomic and qRT-PCR data of plant under different N conditions (Fig. 3, Fig. S4 and 5). Many key genes involved in N transport, assimilation, as well as signaling were found significantly down-regulated in the loss-of-*OsNLP4* mutant and up-regulated in the *OsNLP4*-overexpressing lines. Promoter sequence analysis showed that one or more NRE *cis*-element exist in the promoter of these genes, including *OsNRT2*.*1, OsNRT2*.*2, OsNRT2*.*3, OsNRT2*.*4, OsNRT1*.*1B, OsAMT1*.*1, OsNIA1, OsNIA3* and *OsGRF4*. Through ChIP and Y1H assays, we showed that OsNLP4 directly bound to the promoter fragments of these genes to regulate their expression (Fig. 4). Interestingly, OsNLP4 preferred to regulate the expression of high affinity nitrate/ammonium transporters like *OsNTR2* and *OsAMT1* family members, which may well explain why the *nlp4* mutant phenotype was more severe under low N conditions.

Besides directly regulation, it is worth noting that OsNLP4 can indirectly modulate N utilization through regulating other N signaling regulators. Several regulators, such as OsNRT1.1B, OsGRF4, OsNIGT1 and OsENOD93, were highly expressed in *OsNLP4*-overexpressing rice, while down-regulated in the mutant under HN or LN condition (Fig. 4C). These key regulators had been also proved to significantly improve rice yield and NUE. For example, compared with NRT1.1B-*japonica*, NRT1.1B-*indica* with a single amino acid substitution, results in increased nitrate uptake, root-to-shoot transport and expression of nitrate-responsive genes, and a consequent increase in grain yield and NUE (Hu et al., 2015). Antagonistically working with DELLA, OsGRF4 boost rice yield and NUE by promoting and integrating N assimilation, carbon fixation and growth (Li et al., 2018). In addition, transgenic rice plants overexpressing the OsENOD93-1 gene improve NUE with increased shoot dry biomass and seed yield (Bi et al., 2009).

N and C metabolic processes are closely interrelated and tightly co-regulated (Stitt et al., 2002). NUE is not only dependent on N metabolism, and manipulating C metabolism is useful in some cases to improve NUE (Chardon et al., 2012). In addition to modulating N metabolism, OsNLP4 may also participate in many C metabolism processes, such as C fixation and reductive pentose phosphate cycle as implicated by transcriptomic analyses (Fig. S4, Table S2), indicating that OsNLP4 coordinates N and C assimilation simultaneously.

According to the publicly available transcriptome data (Hruz et al., 2008), rice NLPs are expressed in almost all organs, with OsNLP1 and OsNLP3 preferentially expressed in source organs and very low transcriptional level of *OsNLP6*. Different from the other 5 *OsNLPs*, expression of *OsNLP4* is repressed by many abiotic stresses and induced by low phosphate availability (Chardin et al., 2014). In our study, we found OsNLP4 is widely expressed in all organs, with the highest in leaf and followed by root (Fig. S7). Its expression in root and leaf is rapidly and dramatically induced by nitrogen starvation, and repressed by nitrate replenishment. Ammonium can also depress the expression of *OsNLP4*, but with a weaker suppression than nitrate, suggesting that *OsNLP4* plays a more extensive role in nitrogen metabolism. (Fig. 6A). According to the report of Cao et. al. (2017), low nitrate induces the expression of *ZmNLP3* and *ZmNLP4* in maize. However, all the *NLPs* in Arabidopsis are not significantly induced or repressed in the N-starved seedlings resupplied with nitrate for one hour. Arabidopsis NLP6- and NLP7-regulated N assimilation and signaling is mostly at post-translational level (Konishi and Yanagisawa, 2013; Marchive et al., 2013). The different responds to environmental N between *OsNLP4, ZmNLP3/4* and Arabidopsis *NLPs* imply there may also exist transcriptional regulation in rice and maize *NLPs*. As an important modulator of N utilization, it is reasonable for NLPs to respond at both transcriptional and post-transcriptional levels to maximize N utilization.

Arabidopsis NLP7 has been proved as key transcription factor of nitrate signaling and primary responses through nuclear retention mechanisms (Marchive et al., 2013). Similar as in Arabidopsis, we found OsNLP4 also showed nuclear retention in respond to nitrate. Under nitrate starvation, weak OsNLP4-GFP signals were found in cytosol. However, after nitrate was resupplied, OsNLP4-GFP proteins were quickly and dramatically accumulated in the nucleus in 30 min and moved to cytosol in 1h after nitrate deprivation (Fig. 6C-G). As *OsNLP4* is strongly induced by N-free condition, while less OsNLP4 protein exists without the presence of N. This dis-correlation between transcription and translation implies that *OsNLP4* prepares for N deficiency by responding rapidly at the transcriptional level, priming the *OsNLP4* mRNA capacity and readiness for protein synthesis. Once N is available in the growth environment, *OsNLP4* mRNA can be rapidly translated to promote N absorption and utilization. This may be an economic and effective response mechanism for plants to adapt to the changes of nitrogen sources in the environment.

In conclusion, our results demonstrate that OsNLP4, as a master regulator in coordinating N uptake, assimilation and signaling, can significantly improve the grain yield and NUE under different N conditions, and is a promising candidate gene for improving yield and NUE of the crops.

## Methods

### Plant materials and growth conditions

The rice (*Oryza sativa* L.) varieties Zhonghua 11 (ZH11) and Dongjin (DJ) were used in this study. Two mutants *osnlp4-1* and *osnlp4-2* were obtained by CRISPR technology. The homozygous *osnlp4-3* T-DNA insertion mutant (PFG_1B-00433.L, DJ background) was ordered from the Korea Rice Mutant Center (Pohang, Korea) (Jeong et al., 2002). ACTIN1::OsNLP4 overexpression construct was made by inserting the coding region of OsNLP4 into pCB2006 via GATEWAY cloning system (Lei et al., 2007). For hydroponic culture, rice seedlings were grown in modified Kimura B solution in an artificial climate chamber with a 16-hour light (30°C)/8-hour dark (28°C) photoperiod, ∼300 µmol m^-2^ s^-1^ photon density, and ∼60% humidity. The modified Kimura B solution contained the following macronutrients (mM): (NH_4_)_2_SO_4_ (0.5), MgSO_4_·7H_2_O (0.54), KNO_3_ (1), CaCl_2_ (0.36), K_2_SO_4_ (0.09), KH_2_PO_4_ (0.18), and Na_2_SiO_3_·9H_2_O (1.6); and micronutrients (µM): MnCl_2_·4H_2_O (9.14), H_3_BO_3_ (46.2), (NH_4_)_6_Mo_7_O_24_·4H_2_O (0.08), ZnSO_4_·7H_2_O (0.76), CuSO_4_·5H_2_O (0.32), and Fe(II)-EDTA (40), with the pH adjusted to 5.8. The nutrient solution for culture was renewed every three days. The modified Kimura B solution with different N concentration (20 µM, 0.2 mM and 2mM KNO_3_ (or NH_4_Cl)) were used for hydroponic cultivation.

The *Arabidopsis thaliana* ecotype Columbia (Col-0) was used in this study. Seeds were sterilized with 15% bleach for 12 min, and then washed five times with sterile water. Sterilized seeds stratified at 4 °C for 2 days, and plated on solid medium containing 1% (w/v) sucrose and 0.6% (w/v) agar. Nitrate-less medium was modified on MS medium with KNO_3_ as sole N source: 10 mM nitrate medium (similar to MS except 20 mM KNO_3_ and 20 mM NH_4_NO_3_ was replaced with 10 mM KCl and 10 mM KNO_3_), 3 mM nitrate medium (similar to MS except 20 mM KNO_3_ and 20 mM NH_4_NO_3_ was replaced with 7 mM KCl and 3 mM KNO_3_), 1 mM nitrate medium (similar to MS except 20 mM KNO_3_ and 20 mM NH_4_NO_3_ was replaced with 9 mM KCl and 1 mM KNO_3_). To investigate different growth rates under different N conditions, seeds were germination and grew on medium containing different concentrations of nitrate at 22 °C under 16-h light/8-h dark photoperiod. For evaluation the phenotype of soil-grown plants, seeds were germinated and grew in soil at 22 °C under 16-h light/8-h dark photoperiod.

### RNA extraction and qRT-PCR

Total RNA isolated using Trizol reagent (Invitrogen, Carlsbad, California, USA) were used for reverse transcription. qRT-PCR was performed with a StepOne Plus Real Time PCR System by using a TaKaRa SYBR Premix Ex Taq II reagent kit. Rice *Actin1* was used as the internal reference. All the primers used are shown in Supplementary Table 4.

### Promoter-GUS analyses

A 3.0-kb promoter region of *OsNLP4* was amplified from ZH11 and cloned into pCB308R (Lei et al., 2007; Xiang et al., 1999) to generate *OsNLP4*promoter::GUS, and the resulting vector was transformed into ZH11. For GUS staining, seedlings of *OsNLP4*promoter::GUS transgenic rice under different nitrogen conducts were sampled for histochemical detection of GUS expression. After staining for 12 hours at 37°C, the samples were dehydrated in an ethanol series (70, 85, 95, and 100%) to remove the chlorophyll. The stained tissues were observed under a HiROX MX5040RZ digital optical microscope (Questar China Limited) and photographed using a digital camera (Nikon D700).

### Time-course analyses of OsNLP4 expression in response to N availability

Rice seedlings (ZH11) were cultured in modified Kimura B solution with 0.5 mM KNO_3_ as the sole N source for 10 days. Then, seedlings were cultivated in modified Kimura B solution without nitrogen for 8 hours, and transferred to 5 mM KNO_3_ or NH_4_Cl (5 mM KCl used as the control) in modified Kimura B solution for nitrogen induction. Roots and shoots were collected at 0, 0.5, 1, 2, 4, 6 and 8 h, and the expression of OsNLP4 was determined using qRT-PCR. Growth-chamber conditions were 16-hour light (30°C)/8-hour dark (28°C) photoperiod, ∼300 µmol m^-2^ s^-1^ photon density, and ∼60% humidity.

### Subcellular localization assay

To investigate the subcellular localization of OsNLP4, the ACTIN1::OsNLP4-GFP fusion constructs were produced by inserting the full length CDS of OsNLP4 into the pCB2006-ACT::GFP vector, and the resulting vector was transformed into ZH11. To investigate the nuclear-cytoplasmic shuttling of OsNLP4, rice seedlings were germinated and grown on the modified Kimura B solution without N for 10 days, then treated with 50 mM KNO_3_ for 60 min, and removed them out finally. Laser scanning confocal imaging was performed using the Zeiss 710 microscope equipped with an argon laser (488nm for green fluorescent protein (GFP) excitation).

### Metabolite analyses

The metabolite analyses were performed on the seedlings of 16-day-old plants grown hydroponically with different concentrations of nitrate. The percent total N content in oven-dried plant material was measured with an NC analyzer (Vario EL III model, Elementar, Hanau, Germany) according to the manufacturer’s instructions. Nitrate was extracted in 50 mM HEPES–KOH (pH 7.4), and measured by the method according to Cataldo (Cataldo et al., 2008). The maximum in vitro activity of NR was assayed as described previously (Sylvie et al., 1998).

### Uptake of ^15^N-nitrate

^15^N-accumulation assay after ^15^N-nitrate labeling was performed with ^15^N-labeled KNO_3_ (99 atom % ^15^N, Sigma-Aldrich, no. 335134). For ^15^N-nitrate accumulation assay, rice seedlings were cultured in the Kimura B solution for 10 days. Next, the seedlings were pre-treated with the Kimura B solution for 2 hours and then transferred to modified Kimura B solution containing 5 mM ^15^N-KNO_3_ for 24 hours. At the end of labeling, the roots were washed for 1 minute in 0.1 mM CaSO_4_. Seedlings were then dried at 70 °C to a constant weight and grinded. ^15^N content was analyzed using a continuous-flow isotope ratio mass spectrometer (DELTA V Advantage) coupled with an elemental analyzer (EA-HT, Thermo Fisher Scientific, Inc., Bremen, Germany).

### Chlorate sensitivity assay

Seedlings were first cultured in modified Kimura B solution containing 2 mM KNO_3_ for 4 d after germination. Seedlings were subsequently treated with 2 mM chlorate for 4 d and allowed to recover in modified Kimura B solution (2 mM KNO_3_) for 2 d. Chlorate sensitivity was calculated by the mortality rate of plants poisoned by chlorate.

### Field tests of rice

To investigate the application potential of OsNLP4, field tests for rice using OsNLP4-OE plants and *nlp4* mutants in the ZH11 background (*nlp4-3* was DJ background) were carried out in the paddy field under natural growth condition during 2017-2019 at two experimental stations: Lingshui and Chengdu. For the field test in Chengdu in 2019 (April to September), urea was used as the N source with 80 kg N/hm^2^ for low N, 200 kg N/hm^2^ for normal N and 500 kg N/hm^2^ for high N. The plants were transplanted in 10 rows × 20 plants for each plot (8 m^2^), and four replicates were used for each N condition. For field test in Lingshui (December 2017 to April 2018), urea was used as the N source, and the planting density was 8 rows × 10 plants for each plot (3.4 m^2^) with four replicates under normal N application condition (100 kg N/hm^2^). To reduce the variability in field test, the fertilizers are evenly applied to every plot for each N application level. For the final field test, the edge lines of each plot were removed to avoid margin effects.

### Analysis of yield and NUE

All the agronomic traits were analyzed as described previously (Hu et al., 2015). All filled grains from a single plant were collected and dried at 60 °C in an oven for measurements of grain yield per plant. All grains in a single plot were collected and treated as described above for measurements of actual yield.

NUE was defined as the total amount of yield in the form of grain or biomass achieved per unit of available N (actual yield per plot / total N application per plot) (Hawkesford, 2014).

### RNA-sequencing analysis

Each stain has about 100 seedlings (ZH11 background) of every treatment was grown hydroponically in a growth chamber with the condition described above. The seedlings were cultured in modified Kimura B solution after germination with different nitrate concentrations (0.02/2 mM). 16-day-old seedlings were sampled for RNA-sequencing. For each treatment, 20 seedlings were collected as a sample, and three independent biological replicates were conducted. RNA library construction and sequence analysis were conducted as described previously (Li et al., 2016).

### Chromatin immunoprecipitation

Seedlings were grown in modified Kimura B solution (0.02 mM or 2 mM KNO_3_) for 10 days. The ChIP assay was performed as reported previously (Cai et al., 2014). ACTIN1::NLP4-HA transgenic plants, anti-HA antibodies (Abmart), and salmon sperm DNA/protein A agarose beads (Millipore, USA) were used for ChIP experiment. DNA was purified using phenol/ chloroform (1:1, v/v) and precipitated. The enrichments of DNA fragments were quantified by qPCR using specific primers (Supplementary Table 4). Enriched values were normalized with the level of input DNA.

### Yeast-one-hybrid assay (Y1H)

A cDNA fragment encoding *OsNLP4* was amplified, and inserted into plasmid pAD-GAL4-2.1 (AD) to get AD/*OsNLP4*. The putative NRE from the flanking sequences of N uptake and assimilation genes were ligated into the pHIS2 (BD) vector. The constructed vectors were co-transferred into yeast Y187 competent cells and grew the cells on SD-Trp-Leu medium for 3 days at 30□, then transferred to SD-Trp-Leu-His medium with 10 mM or 20 mM 3-aminotriazole (3-AT, sigma) at different dilutions. The yeasts were incubated at 30□ for 5 days and the extent of yeast growth was determined. The AD and BD empty vector were used for negative control.

### Statistical analysis

Statistically significant differences were computed based on the one-way ANOVA.

## Supporting Information

Figure S1. Identification of *OsNLP4* knockout mutants and overexpressing lines.

Figure S2. Loss-of-function of *OsNLP4* results in severe N deficiency phenotype in DJ background.

Figure S3. *OsNLP4* overexpressing plants increased rice yields in the field.

Figure S4. Differentially expressed genes (DEGs) in the WT, ko and OE under

LN and HN conditions.

Figure S5. Transcription levels of selected genes involved in N metabolism, as revealed by RT-qPCR.

Figure S6. Loss of function of OsNLP4 seriously affects N metabolism in DJ background.

Figure S7. Expression pattern of *OsNLP4 promoter-GUS*.

Figure S8. *OsNLP4* restores the N deficiency phenotype in Arabidopsis *nlp7-1* mutant.

Figure S9. OsNLP4 broadly regulates the genes related to N utilization and signaling in Arabidopsis *nlp7-1* mutant.

Table S1. The FPKM values of the genes involved in N metabolism.

Table S2. The FPKM values of the genes involved in C metabolism.

Table S3. The NRE motifs found in the N related genes.

Table S4. Primers used for PCR.

## Author’s contribution

J.W., L.-H.Y. and C.-B.X. designed the experiments. J.W. performed experiments and data analysis, and wrote the manuscript. Z.-S.Z., J.-Q.X., A.A., Y.S., Y.-J.H., G.-Y.W., L.-Q.S., H.T. and Y.L. contributed to assist in performing part of the experiments. S.-M.W., Q.-S.Z., P.Q., Y.-P.W., S.-G.L., C.-Z.M., G.-Q.Z.., contributed to part of the field experiments, C.-C.C contributed some rice genetic materials used in this project. C.-B.X. and L.-H.Y. supervised the project and revised the manuscript.

## Acknowledgements

This work was supported by grants from the National Key R & D Program of China (grant no. 2016YFD0100701), National Natural Science Foundation of China (grant no. 3157110003), China Postdoctoral Science Foundation (grant no. 2015M580544), and Special fund of China Postdoctoral Science Foundation (2016T90577). The authors thank the Rice T-DNA Insertion Sequence Database (RISD) for providing T-DNA insertion lines used in this study.

---

**Figure 1. OsNLP4 influences plant growth under different N conditions**.

(**A**) Phenotypes of wild-type (WT), *nlp4* mutants (*nlp4-1*/*nlp4-2*) and *OsNLP4*-OE lines (OE-1/OE-9) grown in hydroponic medium with different nitrate concentrations for 25 days. Bar = 4.0 cm.

(**B-D**) Seedling biomass. Total fresh weight (B), shoot fresh weight (C), root fresh weight (D) of seedlings were measured. Values are the mean ± SD of four independent replications each containing 10 plants per genotype. *P* values are from the one-way ANOVA (The letters a, b and c indicate significant differences. P < 0.05). FW, fresh weight.

**Figure 2. Field trials show OsNLP4 acting as a master regulator of NUE**

(**A**) Growth phenotype of representative wild-type, *nlp4* mutants (*nlp4-1*), and *OsNLP4*-OE plants (OE-9) at maturing stage in the field trial at Chengdu in 2019. LN, Low N conditions. NN, Normal N conditions. HN, High N conditions. Bar = 10 cm. (**B-D**) Grain yield per plant (B), actual yield per plot (C), NUE (D) of WT, *nlp4* mutants (*nlp4-1*/*nlp4-2*) and *OsNLP4*-OE (OE-1/OE-9) plants under LN, NN, HN conditions in the field trial at Chengdu (2019). Values are the means ± SD (four replications each containing 18 plants for grain yield per plant, 200 plants for actual yield per plot, NUE) (The letters a, b and c indicate significant differences. P < 0.05).

**Figure 3. Expression profiling analysis of rice seedlings in different nitrate concentrations**.

(**A**) Number of up-regulated and down-regulated genes in WT vs. ko-LN, WT vs. ko-HN, WT vs. OE-LN and WT vs. OE-HN, as revealed by RNA-seq.

(**B**) Venn diagrams analysis of the common and specific genes unique and shared between the different pairwise comparisons.

(**C**) Heatmap of the enriched NUE-related genes in DEGs under LN and HN conditions. Genes involved in nitrogen uptake (including high-affinity nitrate transportation, dual-affinity nitrate transportation, low-affinity nitrate transportation and high-affinity ammonium transportation), assimilation (including nitrate assimilation and ammonium assimilation) and signal transduction were shown. The results were presented as log2 ratios. Only results with a P value < 0.05 and that have been confirmed in three independent experiments were included. A color code was used to visualize the data. LN (left figure): 0.02 mM KNO_3;_ HN (right figure): 2 mM KNO_3_.

**Figure 4. OsNLP4 binds to the promoters of many N-related genes directly**.

(**A-I**) OsNLP4 mediated ChIP–qPCR enrichment (relative to input) of NRE-containing promoter fragments (marked with a dot) from (A) *OsNRT2*.*1*, (B) *OsNRT2*.*2*, (C) *OsNRT2*.*3*, (D) *OsNRT2*.*4*, (E) *OsNRT1*.*1B*, (F) *OsNIA1*, (G) *OsNIA3*,

(H) *OsAMT1*.*1* and (I) *OsGRF4*. Data are mean ± SD (*n* = 3). The letters a, b and c indicate significant differences (P < 0.05).

(**J**) Y1H assay. Empty pAD and pBD were used as negative control. Serial yeast dilutions (1:1, 1:10, 1:100 and 1:1,000) were grown on different SD medium for 5 days.

**Figure 5. OsNLP4 regulates N uptake and assimilation**.

(**A**) Chlorate-sensitivity assay in wild-type, *nlp4* mutants and *OsNLP4*-OE plants treated with 2 mM chlorate. Bar = 1.0 cm.

(**B**) Chlorate sensitivity was calculated by the mortality rate of plants poisoned by chlorate. Values are the mean ± SD of three replications each containing 50 plants per genotype (The letters a, b and c indicate significant differences. P < 0.05).

(**C**) ^15^N accumulation assays in wild-type, *nlp4* mutants and *OsNLP4*-OE plants labeled with ^15^N-nitrate. Values are the mean ± SD of three replications each containing 15 plants per genotype (The letters a, b and c indicate significant differences. P < 0.05).

(**D-F**) Contents of total N (D), and nitrate (E), enzyme activities of NR (F) in the plants grown under different nitrate conditions. 14-day-old seedlings grown in hydroponic medium with different concentrations of nitrate were used for metabolite analyses and enzymatic assays as described in Material and Methods. Values are the mean ± SD of four replications each containing 10 plants per genotype (The letters a, b and c indicate significant differences. P < 0.05). DW, dry weight. FW, fresh weight.

**Figure 6. *OsNLP4* expression is responsive to nitrogen availability**.

(**A**) Time-course analyses of *OsNLP4* expression levels in response to N availability. Plants grown under NN condition were transferred to hydroponic medium without N for 8 h, and then transferred to hydroponic medium with 5 mM KNO_3_, 5 mM NH_4_Cl, or 5 mM KCl). RT-qPCR data are mean ± SD (*n* = 3).

(**B**) GUS staining of 7-day-old *OsNLP7promoter::GUS* transgenic plants cultured under different N conditions. NN: 2 mM KNO_3_; 0N: nitrogen free; N-resupply: nitrogen free for 6 days then resupply with 2 mM KNO_3_ for 24h. Bar = 1.0 cm. (**C-G**) Subcellular localization of OsNLP4. (C) Seedlings grown on nitrogen-free

(**C-G**) medium. (D, E) N-starved seedlings were transferred to a nitrate medium (50 mM) and observed after 15 (D) and 30 min (E). (F) Partial enlargement of figure D in 3 times. (G) N-starved seedlings were transferred to a nitrate medium for 60 min, then to N-free medium again and observed after 60 min. Bar =50 μm.

(**H**) Transcript levels of *OsNLP4* in WT and *OsNLP4* overexpression rice (*ACTIN1::OsNLP4-GFP*) under different N conditions revealed by qRT-PCR. NN: 2 mM KNO_3_; 0N: nitrogen free; re-N: nitrogen free for 9 days then with 2 mM KNO_3_ for 24h. Values are the mean ± SD (*n* = 3). The letters a, b and c indicate significant differences (P < 0.05).

