## supplemental info for "Rice NIN-LIKE PROTEIN 4 is a master regulator of nitrogen use efficiency"

### Supporting information for Rice NIN-LIKE PROTEIN 4 is a master regulator of nitrogen use efficiency by Wu et al.

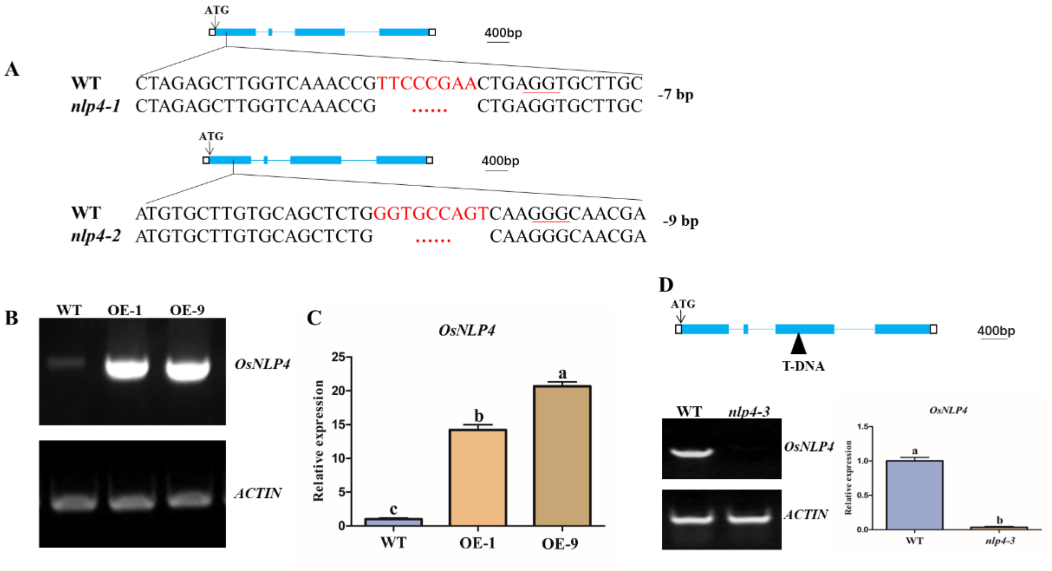


**Figure S1. Identification of *OsNLP4* knockout mutants and overexpressing lines.**

(**A**) Base changes or deletion in *OsNLP4* cDNA of the knockout mutants by CRISPR-cas9-based editing.

(**B**) RT-PCR analysis of *OsNLP4* transcript level in wild type and the OE lines.

(**C**) RT-qPCR analysis of *OsNLP4* expression in wild type and the OE lines.

(**D**) Verification of *nlp4-3* mutant. The triangle represents the T-DNA insertion. RT-qPCR data is mean ± SD ($n$ = 3). The letters a, b and c indicate significant differences. *P* values are from the one-way ANOVA ($P < 0.05$). OsNLP4 gene is illustrated with black-and-white boxes representing the exons and untranslated regions (UTR), respectively, and introns are represented by lines between exons.

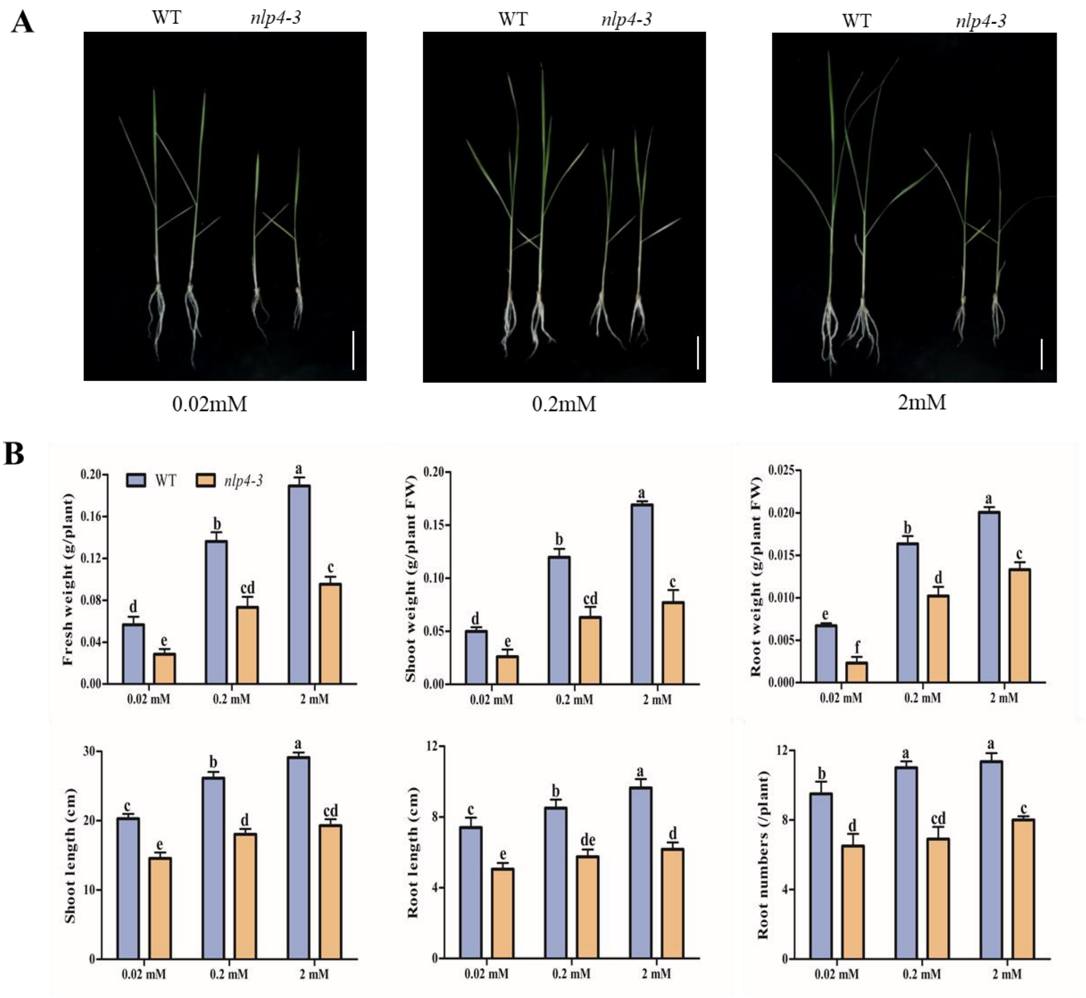


**Figure S2. Loss-of-function of *OsNLP4* results in severe nitrogen deficiency phenotype in DJ background.**

(**A**) Phenotypes of wild-type (WT) and *nlp4-3* mutant grown in hydroponic medium with different nitrate concentrations for 25 days. Bar = 2.0 cm.

(**B**) Measurements of different physiological parameters including the total fresh weight, shoot fresh weight, root fresh weight, root length, root number and shoot length of seedlings. Values are the mean ± SD of four independent replications each containing 10 plants per genotype. *P* values are from the one-way ANOVA (The letters a, b and c indicate significant differences. $P < 0.05$). FW, fresh weight.

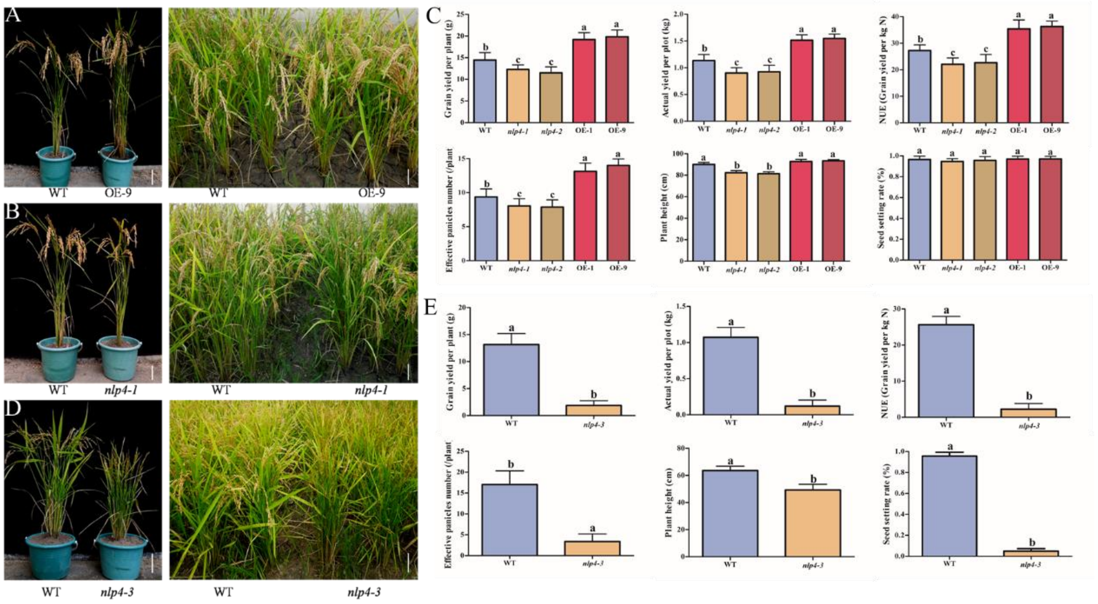


**Figure S3. *OsNLP4* overexpressing plants increased rice yields in the field.**

(**A, B**) Growth of wild-type ZH11, *nlp4-1* and *OsNLP4*-OE plants (OE-9) in the field under normal fertilization level at Lingshui in 2017. Bar = 10 cm.

(**C**) Grain yield per plant, actual yield per plot, NUE, effective panicles number, plant height and seed setting rate of WT, *nlp4* mutants (*nlp4-1*/*nlp4-2*) and *OsNLP4*-OE (OE-1/OE-9) plants under normal nitrogen condition in the field trial at Lingshui (2017). Values are the means ± SD (four replications each containing 30 plants for Plant height and panicles number, 15 plants for seed setting rate and grain yield per plant, 80 plants for actual yield per plot and NUE) (The letters a, b and c indicate significant differences. $P < 0.05$).

(**D**) Growth of wild-type DJ and *nlp4-3* in the field at normal fertilization level at Lingshui in 2017. Bar = 10 cm.

(**E**) Grain yield per plant, actual yield per plot, NUE, effective panicles number, plant height and seed setting rate of WT and *nlp4-3* plants under normal nitrogen condition in the field trial at Lingshui (2017). Values are the means ± SD (four replications each containing 30 plants for Plant height and panicles number, 15 plants for seed setting rate and grain yield per plant, 80 plants for actual yield per plot and NUE) (The letters a, b and c indicate significant differences. $P < 0.05$).

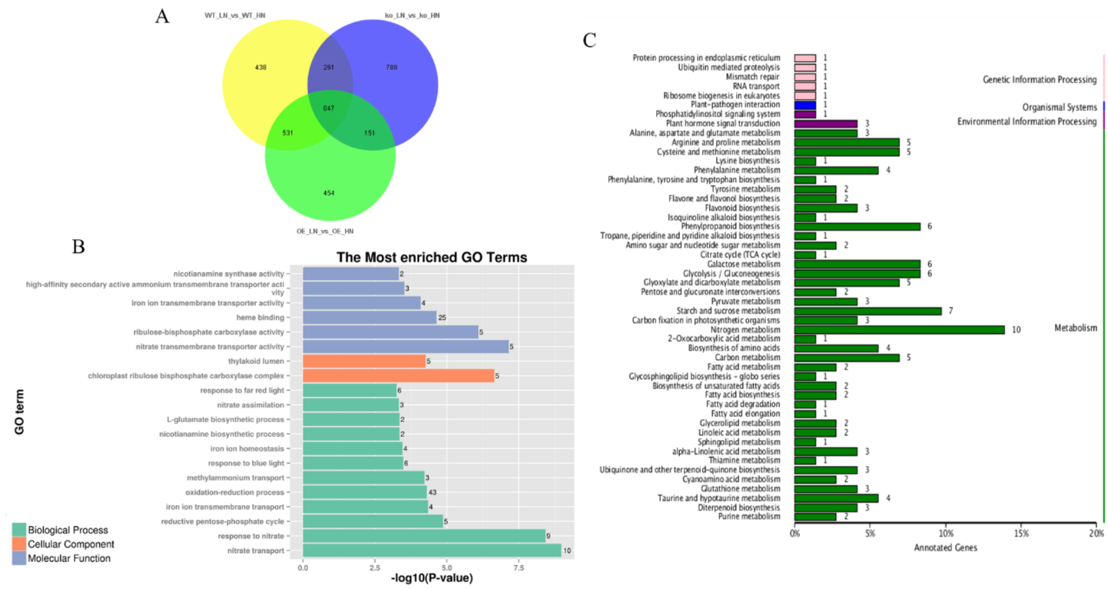


**Figure S4. Differentially expressed genes (DEGs) in the WT, ko and OE under LN and HN conditions.**

(**A**) Venn diagrams analysis of the genes unique and shared between the different pairwise comparisons.

(**B**) Comparative GO (gene ontology) enrichment analysis of enriched DEGs for LN condition (WT vs. ko-LN). The most significantly enriched functional classes of genes were those supporting nitrate transport and response, reductive pentose-phosphate cycle, iron ion transmembrane transport, oxidation-reduction process, methylammonium transport, blue light and far red light response, nicotianamine and L-glutamate biosynthetic process.

(**C**) KEGG pathway analysis of the DEGs for HN condition (WT vs. ko-HN). The greatest number of enriched genes was observed in five pathways: Nitrogen metabolism, Starch and sucrose metabolism, glycolysis / gluconeogensis, galactose metabolism and phenylpropanoid biosynthesis.

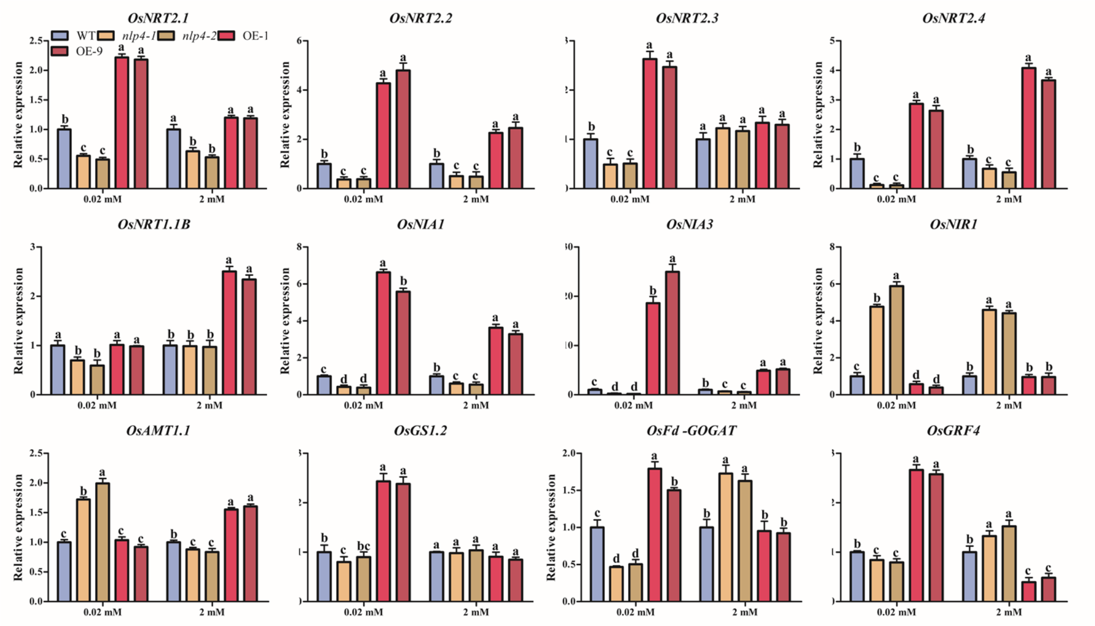


**Figure S5. Transcription levels of selected genes involved in N metabolism, as revealed by RT-qPCR.** 16-day-old seedlings cultured with different nitrate concentrations were sampled for RNA. RT-qPCR data are mean ± SD (*n* = 3). *P* values are from the one-way ANOVA (The letters a, b and c indicate significant differences. P < 0.05).

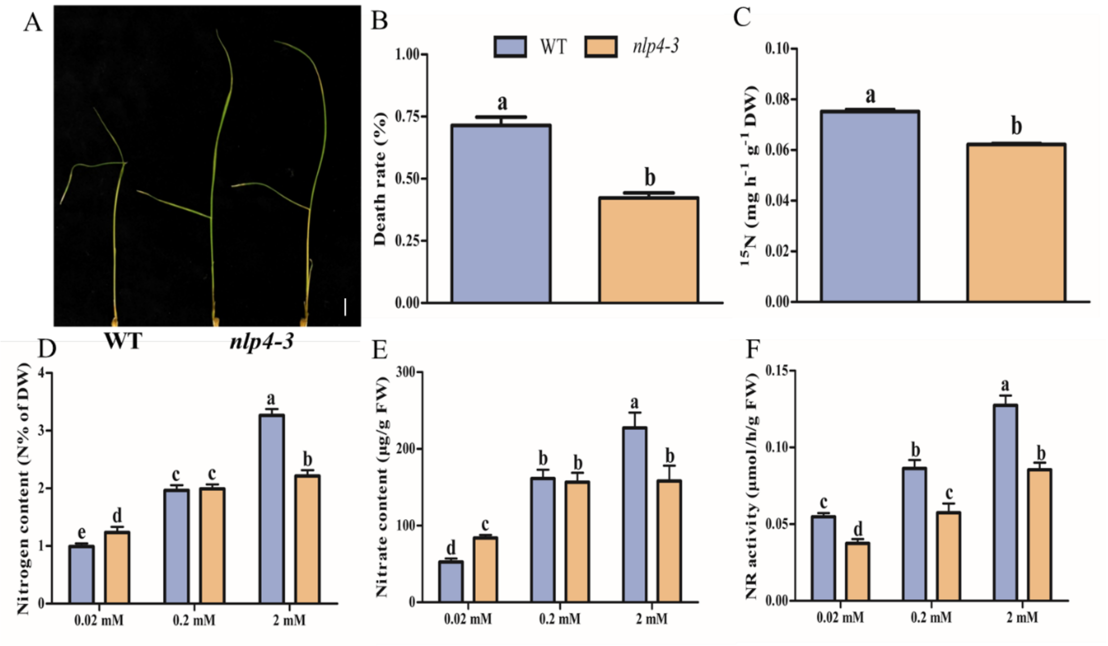


**Figure S6. Loss of function of OsNLP4 seriously affects N metabolism in DJ background.**

(**A**) Chlorate-sensitivity assay in wild-type and *nlp4-3* treated with 2 mM chlorate. Bar = 1.0 cm.

(**B**) Chlorate sensitivity was calculated by the mortality rate of plants poisoned by chlorate. Values are the mean ± SD of three replications each containing 50 plants per genotype (The letters a, b and c indicate significant differences. $P < 0.05$). DW, dry weight. FW, fresh weight.

(**C**) $^{15}$N accumulation assays in wild-type and *nlp4-3* labeled with $^{15}$N-nitrate. Values are the mean ± SD of three replications each containing 15 plants per genotype (The letters a, b and c indicate significant differences. $P < 0.05$).

(**D-F**) 14-day-old seedlings grown in hydroponic medium with different concentrations of nitrate were used for metabolite analyses and enzymatic assays as described in Material and Methods. (D, E) Contents of total nitrogen (D), and nitrate (E) in the plants grown under 0.02 mM, 0.2 mM and 2 mM nitrate conditions. (F) Enzyme activities of nitrate reductase in the plants under different nitrate conditions. Values are the mean ± SD of four replications each containing 10 plants per genotype (The letters a, b and c indicate significant differences. $P < 0.05$).

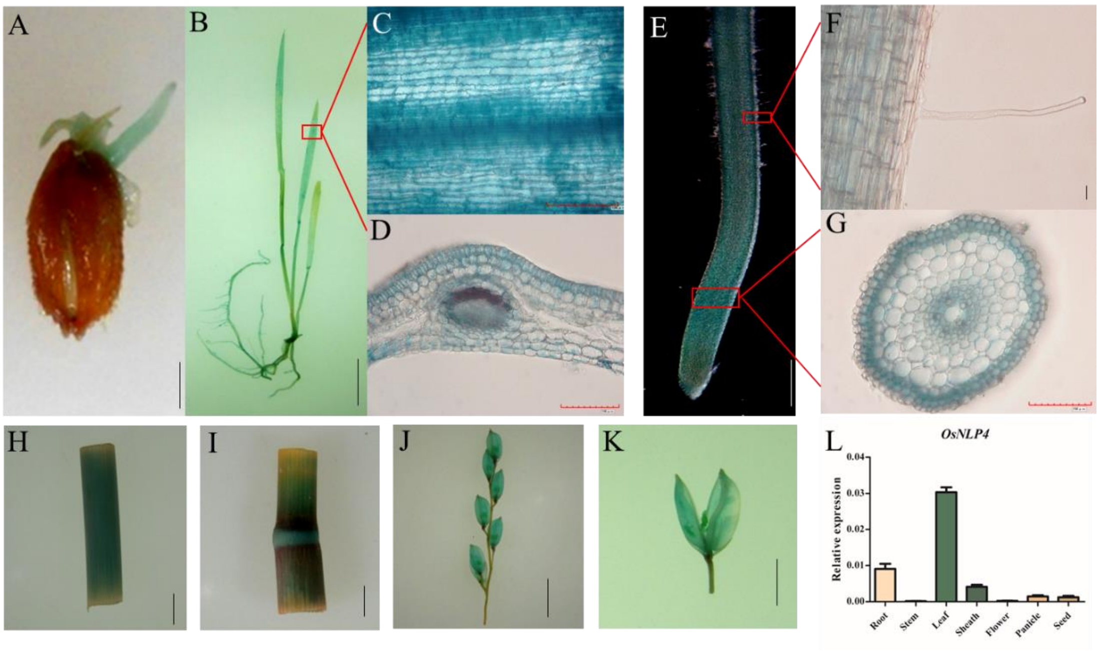


**Figure S7. Expression pattern of *OsNLP4 promoter-GUS*.**

Materials used for GUS staining include germinating seeds (A), 7-day-old seedlings (B), leaf blades (C-D), roots and root hairs (E-G), leaf sheaths (H), stems (I), panicles (J) and caryopses (K). D and G indicate the cross sections of leaf blades and roots respectively. Scale bars, 0.5 cm in A, 2 cm in B, 100 μm in C, D and G, 0.1 cm in E, 10 μm in F, 1 cm in H-J, 0.5 cm in K. (L) Transcription levels of OsNLP4 in different tissues of wild type ZH11. Flowering rice plants were removed from the soil and the roots were washed. RNA was extracted from roots, stems, leaves, leaf sheaths, flowers, flowering panicles and seeds by Trizol. RT-qPCR data are mean ± SD ($n$ = 3).

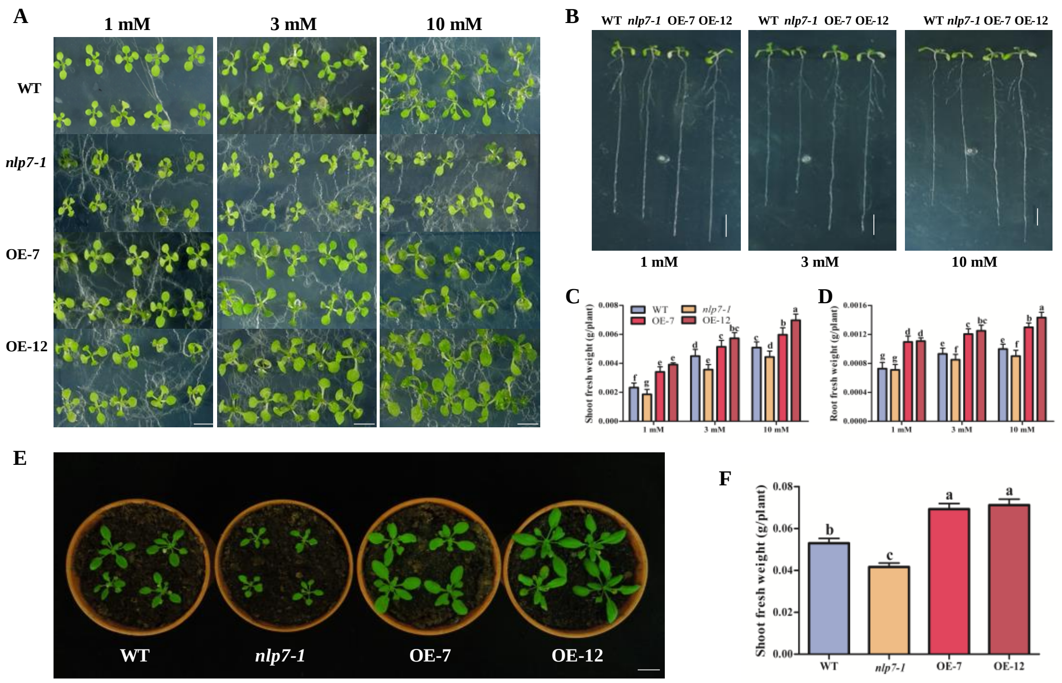


**Figure S8. *OsNLP4* restores the N deficiency phenotype of Arabidopsis *nlp7-1* mutant.**

(**A**) Phenotypes of wild-type (WT), *nlp7-1* and OsNLP4 complementary lines (OE-7/OE-12) grown on medium with different nitrate concentrations for 10 days. Bar = 1.5 cm.

(**B**) Phenotypes of the 14-day-old plants on vertical plates containing different concentrations of nitrate. Bar = 1.0 cm.

(**C, D**) Shoot fresh weight (C), root fresh weight (D) of the plants under different nitrate conditions. Values are the mean ± SD of six independent replications each containing 5 plants per genotype (The letters a, b and c indicate significant differences. $P < 0.05$).

(**E**) Image of 3-week-old OsNLP4 complementary lines, *nlp7-1* and WT plants grew in soil. Bar = 1.0 cm.

(**F**) Shoot fresh weight of the plants grew in soil. Values are the mean ± SD of six independent replications each containing 6 plants per genotype (The letters a, b and c indicate significant differences. $P < 0.05$).

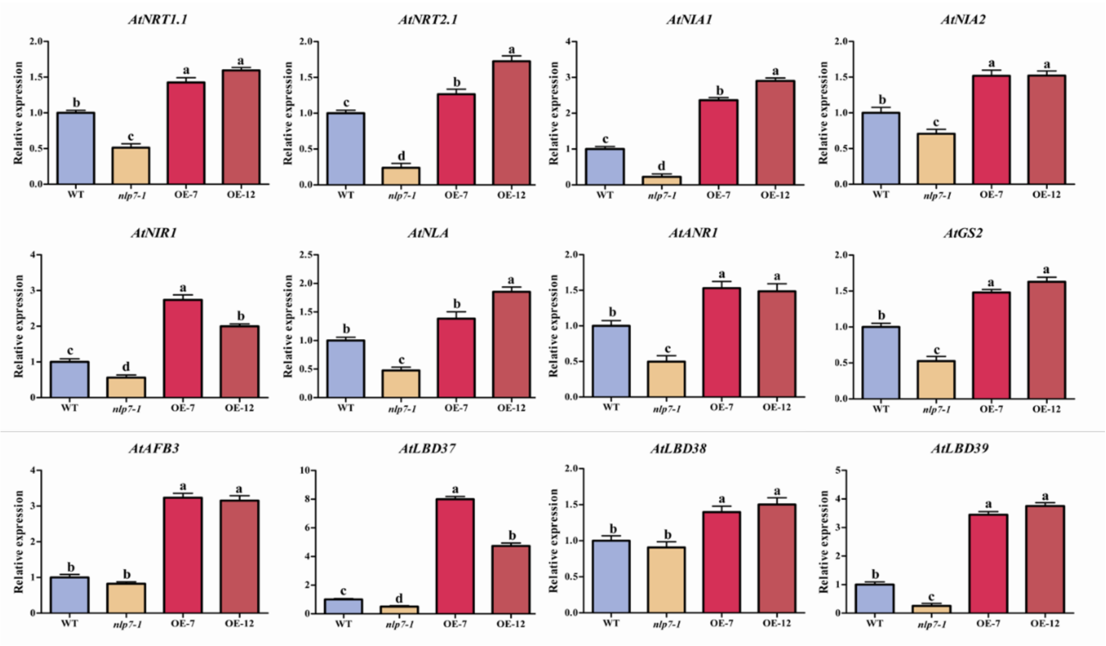


**Figure S9. OsNLP4 broadly regulates the genes related to N utilization and signaling in Arabidopsis *nlp7-1* mutant.** 14-day-old plants grown on MS medium were harvested for qRT-PCR analysis. *UBQ5* was used as an internal control. NRT, nitrate transporter; NIA, nitrate reductase; NIR1, nitrite reductase 1; NLA, nitrogen limitation adaptation; GS2, glutamine synthetase 2; LBD, lateral organ boundary domain; AFB3, auxin signaling F-box 3. Values are the mean ± SD of three replications (The letters a, b and c indicate significant differences. $P < 0.05$).
